# Chromosome-level phased genome assembly of the argan tree *Sideroxylon spinosum* L

**DOI:** 10.1101/2025.04.06.647421

**Authors:** Ivan D. Mateus, Abdellatif Essahibi, Pamela Nicholson, Mohamed Hijri, Ahmed Qaddoury, Laurent Falquet, Didier Reinhardt

## Abstract

Argan (*Sideroxylon spinosum* L. previously known as *Argania spinosa*) is an endemic tree of Morocco, primarily used for seed oil extraction. Increasing interest in its biology and in genes relevant for oil quality and stress resistance demands for high-quality models of its genome and transcriptome. We integrated sequencing data from PacBio HiFi long-reads and Illumina Hi-C to generate independently assembled phased genome models for both parental haplotigs with lengths of 636Mb and 655 Mb, respectively, and with a BUSCO completeness score of >97.8%. Both haplotigs consist of 11 independently assembled fully resolved telomere-to-telomere chromosomes, similar to other species of the Sapotaceae (n = 10-13), and they both contain approximately ∼60% of highly repetitive regions. Comparative analysis with other Sapotaceous genomes suggests overall chromosome conservation, with repeat expansion and chromosome fusion for the two largest chromosomes (chr1, chr2). We also independently assembled the chloroplast genome. This high-quality genome assembly provides a valuable resource for advancing future research on argan biology, its genetic diversity, and traits relevant for fitness and oil production.

## Background & Summary

Argan is a member of the Sapotaceae family (order Ericales), which comprises five tribes and approximately 1,250 species. This family includes economically important species valued for their oil, such as the argan tree (*Sideroxylon spinosum* L., formerly *Argania spinosa* (L.) Skeels), the shea tree (*Vitellaria paradoxa* C.F. Gaertn.), and the miracle fruit tree (*Synsepalum dulcificum* (Schumach. & Thonn.) Daniell). Argan seeds contain an edible oil extensively used for both culinary and cosmetic applications. Argan oil represents a crucial economic resource for many Moroccan families (Lybbert et al., 2011), particularly for women, who play a central role along the entire value chain. Argan is a diploid species with a reported haploid chromosome number ranging between 10 and 12 (El Boukhari et al., 2023).

Argan has not been selectively bred or extensively cultivated, instead, it grows in its natural habitat, where it has been sustainably utilized for centuries. Nevertheless, overgrazing by goats, the intensification of land use in Morocco, and the high demand for argan oil have led to unsustainable exploitation, putting the species at risk (Lybbert et al., 2011). In recent years, rising temperatures, prolonged droughts, and an extended dry season have further exacerbated the challenges for the argan-growing region, known as the “Arganeraie”, which has been listed as UNESCO biosphere heritage in 1998 (Chakhchar et al., 2017).

The economic importance of argan has raised the demand for high-quality genomic resources. A first genome draft containing 75,327 scaffolds (Khayi et al., 2018), was followed by a reference annotation with 62,590 genes (Rupp et al., 2024). The complete mitochondrial and chloroplast genomes have also been released in recent years (Azami et al., 2024; Khayi et al., 2020). Additionally, single-sequence repeat markers (Rabeh et al., 2024) and AFLP markers (Pakhrou et al., 2016) have been developed for the use in population genetics studies, revealing high genetic diversity between populations but low diversity within populations. However, a high-quality genome sequence is required for a more comprehensive understanding of argan evolution, physiology, population genomics, as well as for marker-assisted breeding.

In this study, we used PacBio HiFi sequencing combined with Illumina Hi-C to generate a complete genome assembly of the argan tree with two independently assembled haplotigs and the chloroplast genome. We compared our phased assembly with genome assemblies of two other Sapotaceae species to investigate chromosome composition and evolution. Our telomere-to-telomere genome assembly provides a high-quality reference for comparative genomics, offering new insights into chromosome evolution in this species.

## Methods

### Sample collection and sequencing

Leaf samples of *Sideroxylon spinosum* (syn. *Argania spinosa*) were collected from a specimen at the northern border of the UNESCO biosphere reserve “l’Arganeraie”, along the National Road 207 east of Essaouira at coordinates 31°32’47.9” N 9°21’56.6” W (**Fig. S1**).

We employed a hybrid sequencing strategy that combined PacBio long-read sequencing with Hi-C chromatin conformation data. Genomic DNA was extracted from leaves using a modified CTAB protocol (**Supplementary Methods**). For Hi-C sequencing, nuclei were isolated using a Sucrose/Percoll gradient centrifugation protocol (**Supplementary Methods**). To generate highly accurate long reads, we performed Pacific Biosciences (PacBio) Sequel II HiFi sequencing on sheared genomic DNA (gDNA). Hi-C sequencing was carried out using Illumina NovaSeq 6000 (paired-end, 150 bp) on crosslinked chromatin, which was enzymatically digested with DpnII (^GATC) and HinfI (G^ANTC) using the Arima Genomics Hi-C kit (Cat. #A510008), followed by proximity ligation to capture three-dimensional chromosomal interactions.

### Genome assembly

The genome of *S. spinosum* was assembled by integrating long-read PacBio HiFi sequencing with Hi-C data for physical linkage. High sequence coverage allowed for the independent assembly of the two parental genomes, each consisting of 11 chromosomes. PacBio HiFi sequencing reads were quality-checked using FastQC v.0.11.9 and cleaned with fastp v.0.19.5 (Chen, 2023). The cleaned HiFi reads were assembled into phased haplotigs using hifiasm v.0.16.1 with –hom-cov 50 parameters (Cheng et al., 2021). A K-mer analysis performed with GenomeScope 2.0 (http://genomescope.org) (Ranallo-Benavidez et al., 2020) revealed a low error rate and a high homozygosity between the two haplotigs **(Fig. 1a)**. The haplotigs were further scaffolded using Illumina Hi-C data. We applied HiCup to map the HiC reads to the hifiasm assemblies (Wingett et al., 2015), and we used Juicer v1.6 (Durand, Shamim, et al., 2016) and YAHS v1.2.2 (Zhou et al., 2023) to generate the final telomere-to-telomere (T2T) chromosome assemblies. The YAHS file “telo.c” was customized to include the telomeric sequences of *S. spinosum* TTTAGGG & GGGTGGG and the software was recompiled before generating the final telomer-to-telomer (T2T) chromosome assemblies. The final assembly was manually curated with JuiceBox (v2.15) (Durand, Robinson, et al., 2016) **(Fig. 1b,c)**. The telomeric regions were identified using tidk (v0.2.31) (Brown et al., 2025) with the canonical telomeric sequences TTTAGGG and GGGTGGG (**Suppl. Fig S2**).

**Figure 1.**
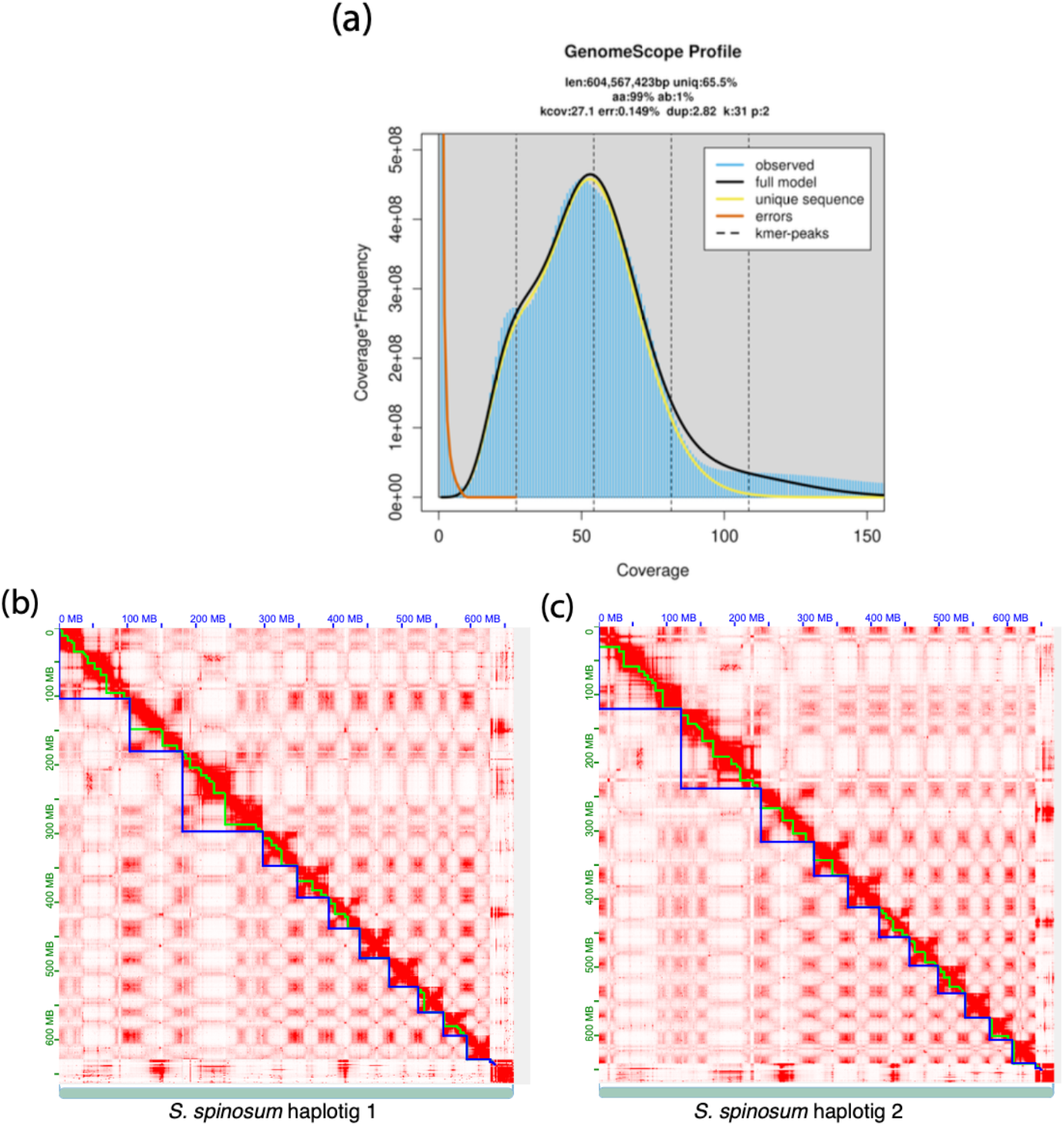
Global genome characterization for *S. spinosum*. **(a)** GenomeScope genome size estimates for *S. spinosum*. **(b,c)** Genome-wide chromosomal HiC maps of haplotig 1 **(b)** and haplotig 2 **(c)** of *S. spinosum*; chromosomes are delineated in blue, and contigs are highlighted with a green outline.

We employed Nucmer (Marçais et al., 2018) and dot (https://github.com/marianattestad/dot) to compare the two parental argan genomes (haplotig 1 and haplotig 2). This comparison revealed extensive synteny and collinearity, with the exception of an inversion on chromosome 2 (**Fig. 2**). Taken together, these analyses confirm the genome structure of 11 chromosomes, as independently validated by the two parental genomes.

**Figure 2.**
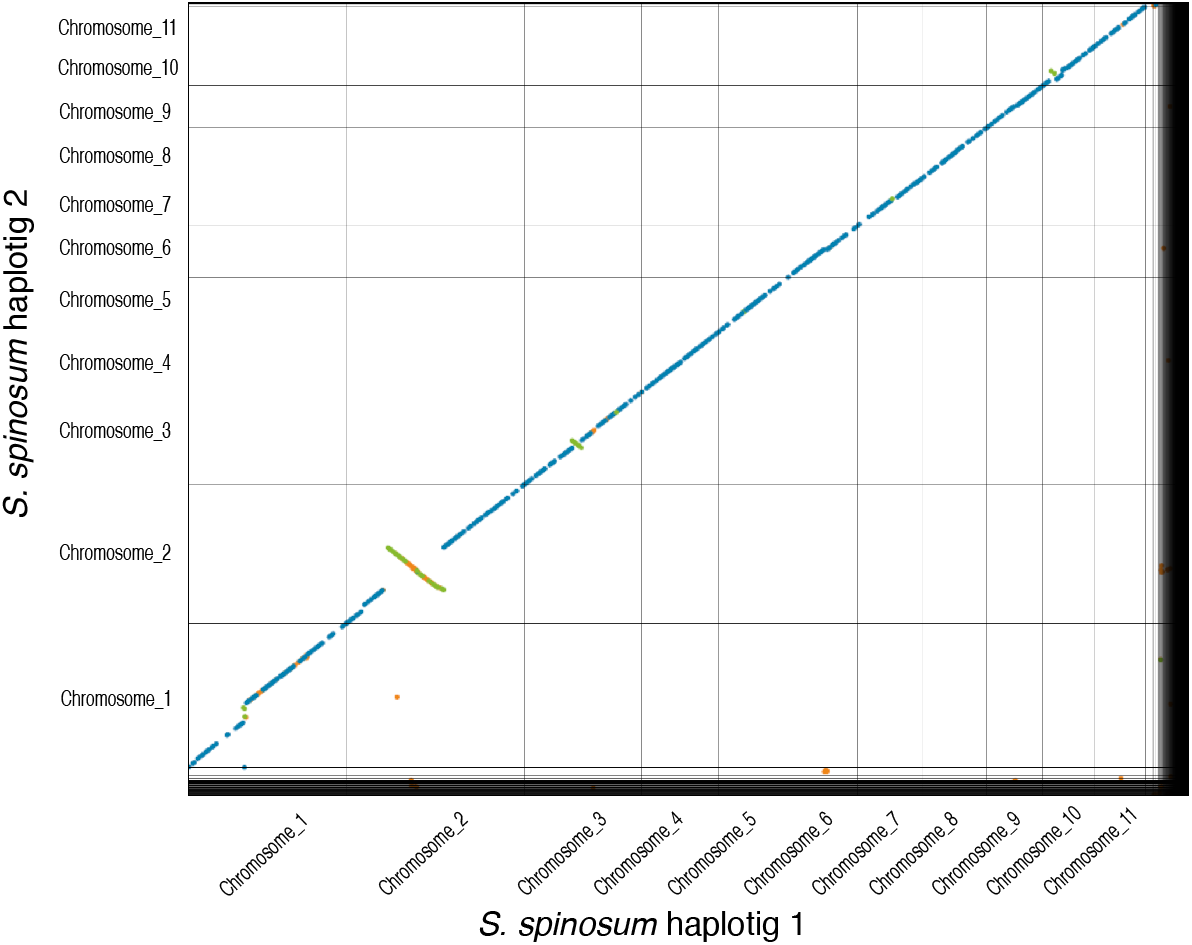
Collinearity between haplotig 1 and haplotig 2 of *S. spinosum*. All chromosomes were highly collinear, except for chromosome 2 that exhibited an inversion.

Most chromosomes contained a strong telomere signal at both ends, with few exceptions (**Fig. S2**). We used Chromsyn (Edwards et al., 2022) to evaluate the synteny between the two haplotigs and a previously published genome assembly of the species (QLOD00000000.2 from PRJNA294096) (Khayi et al., 2018). We observed overall strong synteny between haplotig 1 and haplotig 2 with few inversions near centromeric regions of the larger chromosomes (**Fig. 3**). We also observed that the largest chromosomes (chr1 and chr2) were fragmented in the previously released argan assembly (QLOD) (**Fig. 3**). Evaluation and comparison of the genome assemblies with Quast (Gurevich et al., 2013) revealed that all assembly statistics were improved in our two haplotigs relative to the QLOD assembly (**Supplementary Table 1**), indicating that our new assembly represents a significant upgrade.

**Figure 3.**
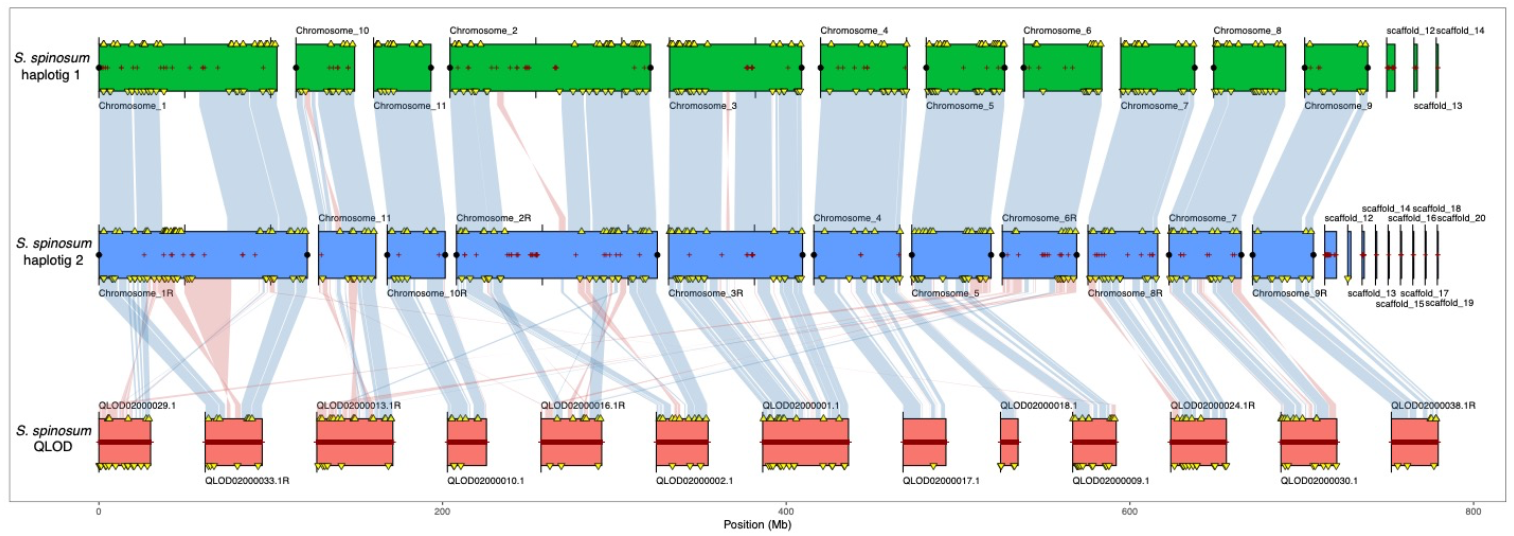
Synteny between different argan genome assemblies. Haplotig 1 and haplotig 2 generated in this study were tested for synteny with the previously published genome assembly QLOD2. Red crosses along chromosomes represent assembly gaps, while black points at the end of chromosomes indicate telomere sequences. Yellow triangles indicate duplicated BUSCO genes in haplotig 1 (n=132) and haplotig 2 (n=138). Blue links between assemblies represent collinear regions. Red links between assemblies represent genome inversions.

The comparison with two other published genomes from the Sapotaceae family revealed similar chromosome numbers. The shea tree (*Vitellaria paradoxa*) contains 12 chromosomes (Hale et al., 2021) and the miracle fruit tree (*Synsepalum dulcificum*) contains 13 chromosomes (Yang et al., 2022). The genomes exhibited extensive synteny and collinearity along the chromosomes, though they displayed multiple chromosomal rearrangements and inversions (**Fig. 4**). Interestingly, the larger chromosomes in argan (chromosome 1 and chromosome 2) appear to have emerged as a result of repeat expansions and chromosome fusions. However, based on the available data, it is unknown which configuration (split or fused chromosomes) represents the ancestral state.

**Fig. 4.**
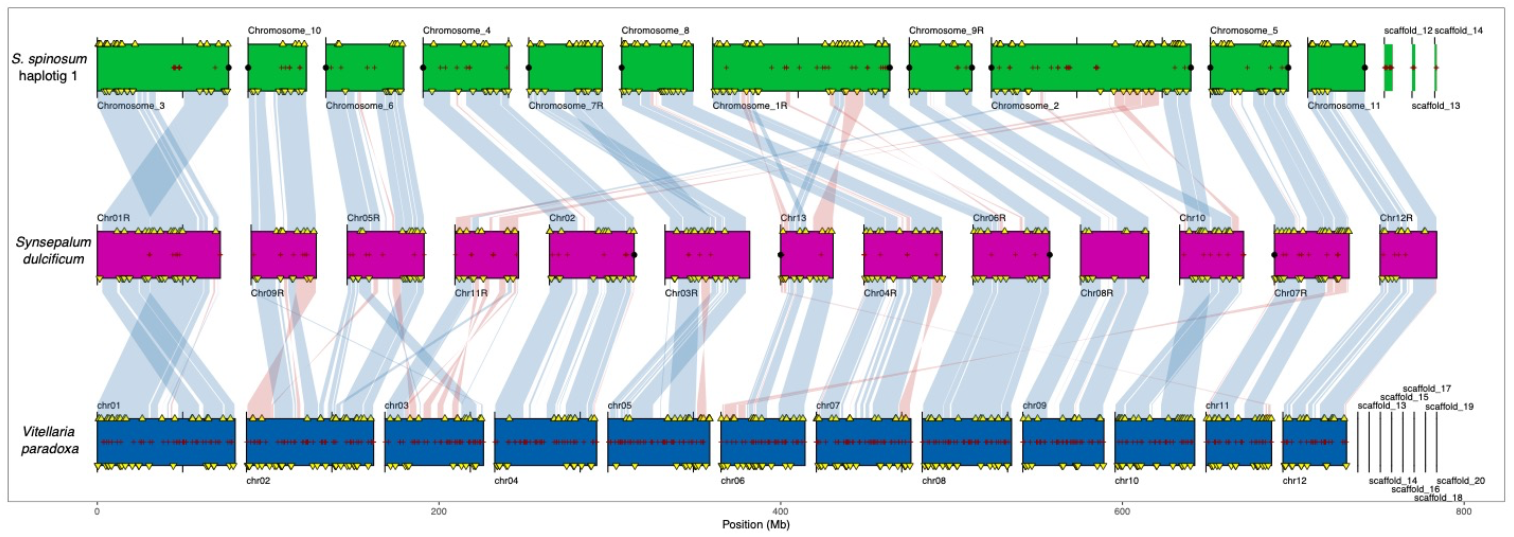
Synteny between three Sapotaceae species. The chromosomal complement of *S. spinosum* (green), *S. dulcificum* (magenta) and *V. paradoxa* (blue) were aligned. For clarity, only the *V. paradoxa* contigs exceeding 1M bp are shown. Red crosses along the chromosomes represent assembly gaps, while black points at the ends of chromosomes represent telomere sequences. Yellow triangles indicate duplicated BUSCO genes. Blue links between assemblies represent collinear regions. Red links between assemblies represent genome inversions.

### Repeat prediction

Repeat elements (including transposable elements) were predicted with RepeatModeler v2.0.1 (Flynn et al., 2020), followed by soft-masking the genome assemblies with RepeatMasker v4.1.2 (Smith et al., 2010). We found that 60.86% of the hap1 and 60.78% of the hap2 genome consisted of repetitive regions (**Supplementary Table 2**). Additionally, the two longest chromosomes in both hap1 and hap2 genomes displayed large expansions of repetitive sequences, each encompassing more than 30 Mb (**Fig. 5 a, b**).

**Figure 5.**
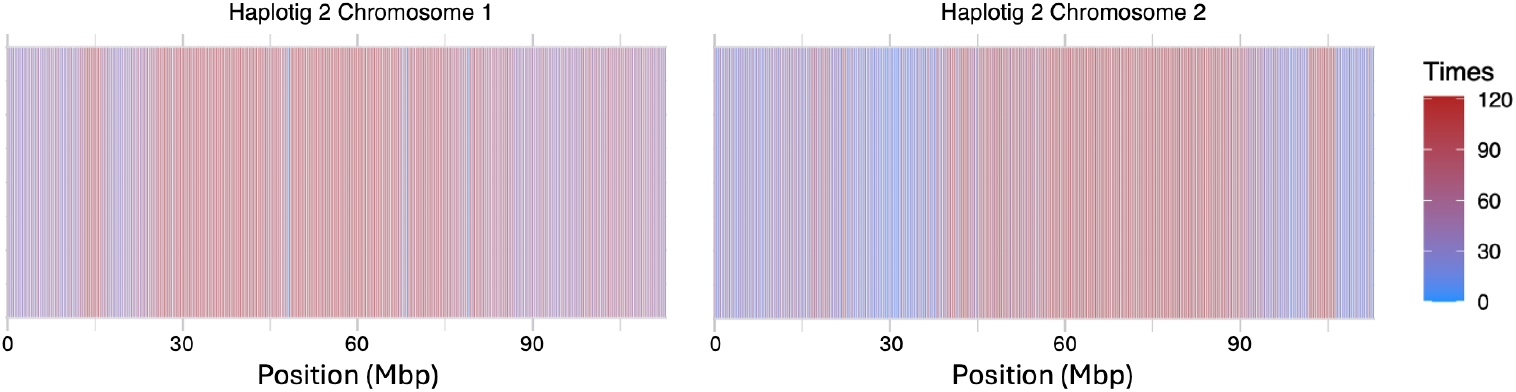
Global organization of the two largest chromosomes (chr1, chr2). Density of predicted protein-coding genes (blue) and repeats (red) along chromosome 1 (left) and chromosome 2 (right) from haplotig 2.

### Chloroplast genome assembly

An independent approach was used to assemble the chloroplast genome. First, the HiFi reads were mapped to the published chloroplast reference genome of *S. spinosum* (MK533159.1) (Khayi et al., 2020) with minimap2 (Li, 2018). The mapped reads were then recovered with SAMTools (Li et al., 2009), and the chloroplast genome was *de novo* assembled from the recovered reads using flye v.2.9.5 (Kolmogorov et al., 2019) with parameters --meta –pacbio -g 150k. We then used Bandage v.0.8.1 (Wick et al., 2015) to identify sequences that display circularity features. Finally, the assembly was annotated and visualized with the Proksee online tool (Grant et al., 2023), which included Prokka annotation (Seemann, 2014). Collinearity between our assembly and published assemblies were evaluated with D-genies (Cabanettes & Klopp, 2018).

The chloroplast sequence consisted of 132,913 bp assembled from 3 contigs with 131 annotated features (**Fig. 6a**). Our new chloroplast assembly was blasted to a previously published chloroplast reference genome resulting in complete blast homology with excellent collinearity. The only difference with the published chloroplast sequence was a 26 kb duplication on the reference chloroplast assembly (**Fig. 6b**).

**Figure 6.**
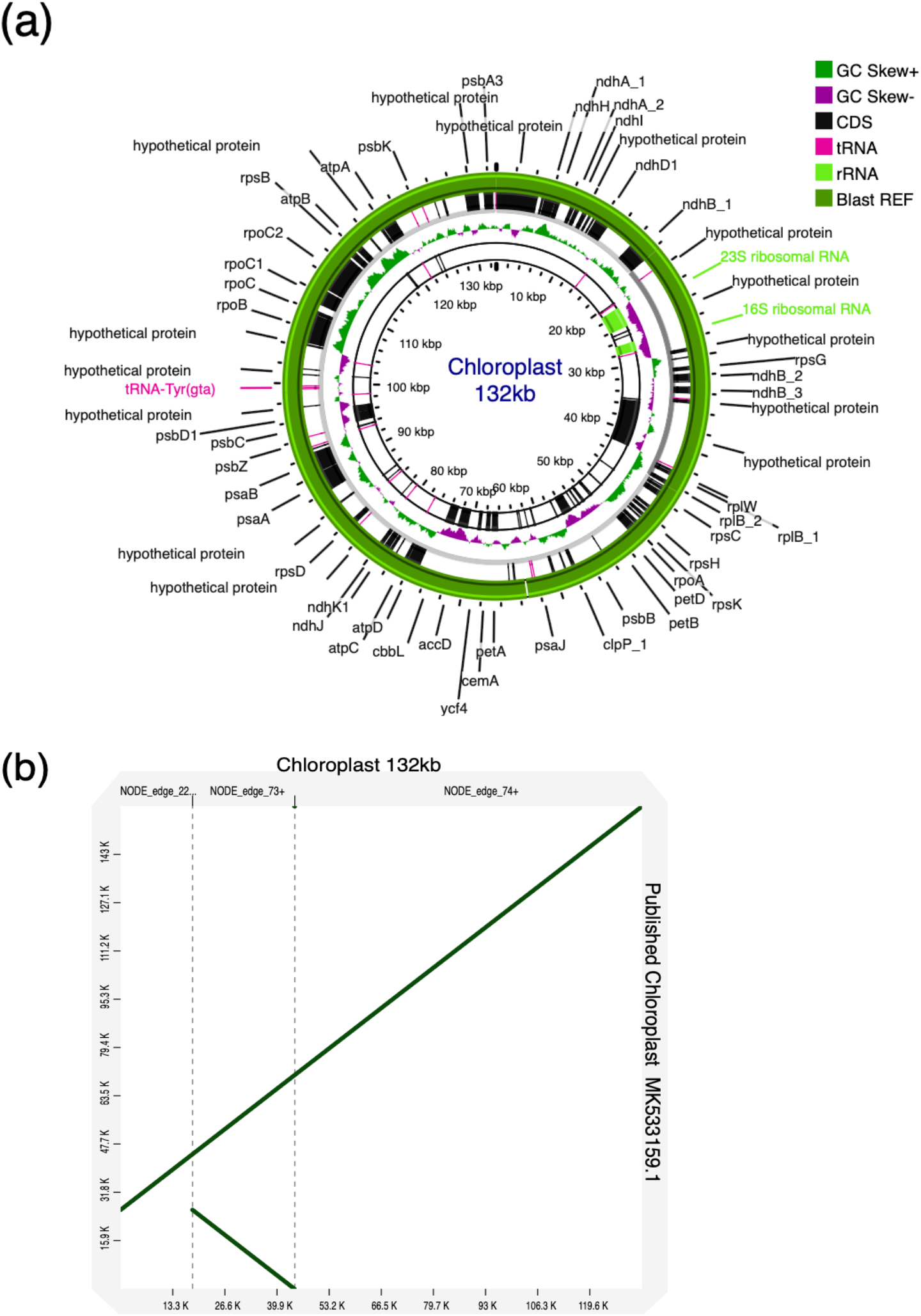
Chloroplast *denovo* assembly. **(a)** Annotated circular visualization of the assembled chloroplast. **(b)** Dot-plot between the newly assembled chloroplast genome and the published chloroplast genome.

### Technical Validation

Genome completeness was assessed using BUSCO v5.4.2 (Manni et al., 2021) with the eudicots_odb10 plant database, which contains 2,326 BUSCO entries. We found that 97.8% and 98.5% for hap1 and hap2, respectively, consisted of complete single-copy BUSCO (**Supplementary Tables 3, 4**).

### Data records

The three PacBio HiFi runs, and the Illumina Hi-C data can be accessed at the ENA repository PRJEB59883 using the identification numbers ERR10968128, ERR10969898, ERR13030747, and ERR13030748, respectively. The genome assembly result is available in the ENA repository numbers PRJEB88017 (hap1) and PRJEB88018 (hap2).

## Code availability

The genome assembly scripts are available in https://github.com/ivandamg/Argan_Genome. All commands and pipelines used in data processing were executed according to the manual and protocols of the corresponding bioinformatic software.

## Author contributions

L.F., I.D.M. and D.R. contributed to the research design. A.E. and M.H. collected the samples. I.D.M., L.F. and P.N. analyzed the data. L.F., D.R., I.D.M. and A.Q. wrote the draft manuscript and revised the manuscript. All co-authors contributed to this manuscript and approved it.

## Additional information

Correspondence and requests for materials should be addressed to L.F. and D.R.

## Acknowledgements

We would like to thank the Next Generation Sequencing Platform of the University of Bern for performing the high-throughput sequencing experiments for this difficult project.

