## Supplemental Material for "Chromosome-level phased genome assembly of the argan tree *Sideroxylon spinosum* L"

### Supplementary Information

**Figure S1**

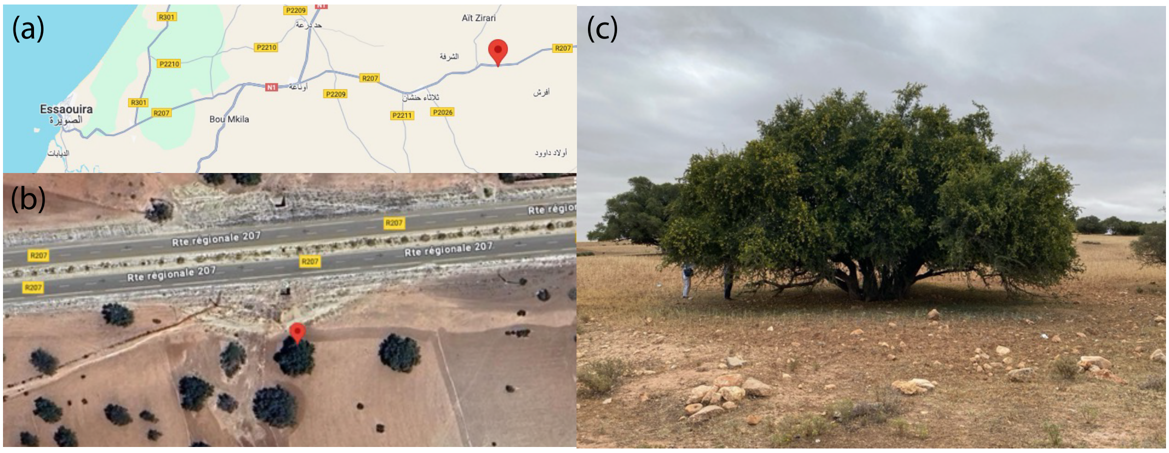


**Figure S1**. Location of the argan tree sampled for genome sequencing. **(a)** and **(b)** show the exact location of the argan tree that was sampled for genome sequencing, while **(c)** displays a photograph of the tree taken during sampling (coordinates 31°32'47.9" N 9°21'56.6" W).

**Figure S2**

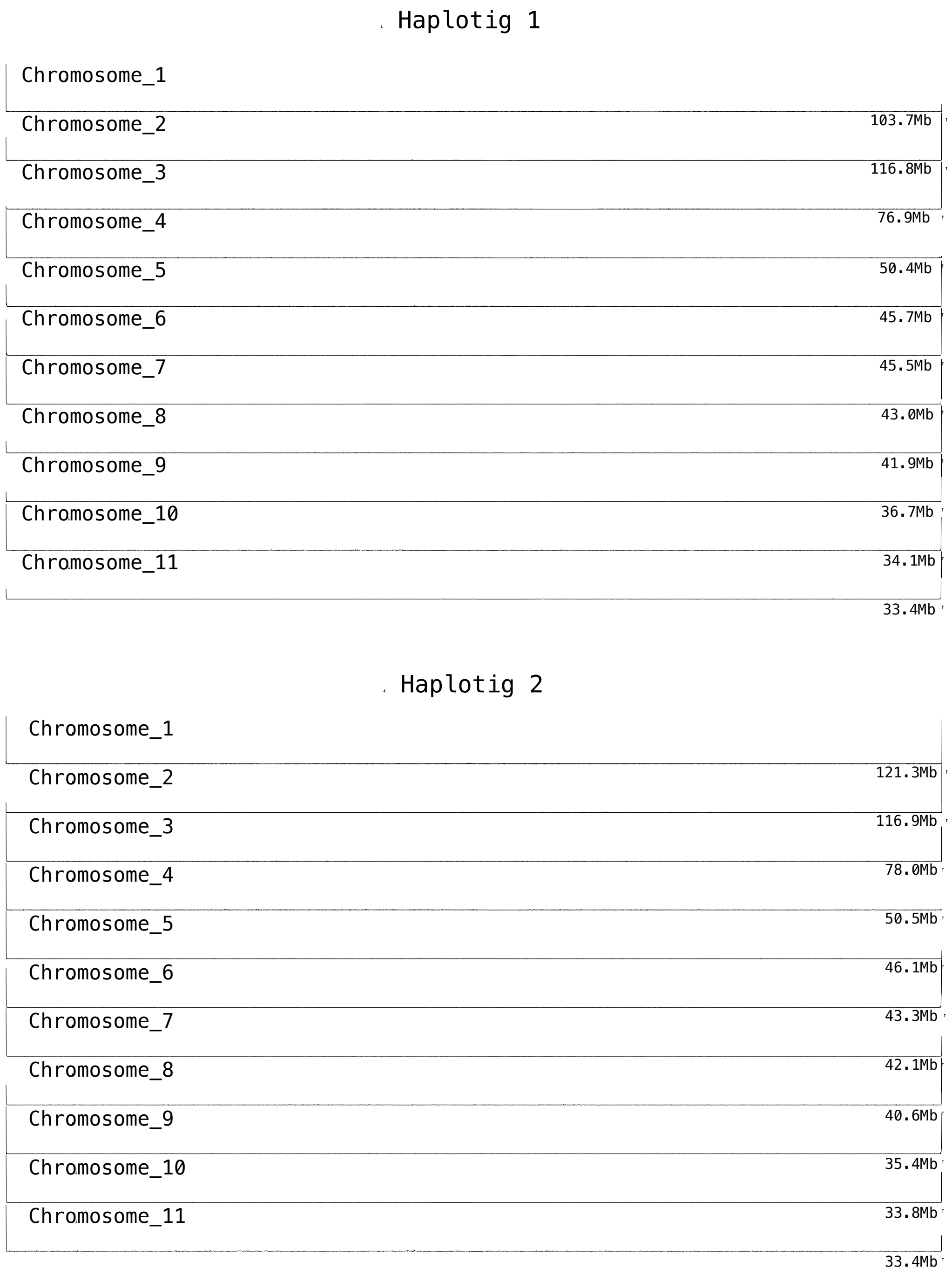


**Figure S2**. Tidk plot representing the telomeres (vertical lines) of haplotig 1 and haplotig 2 at the 5’ and 3’ ends of chromosomes (horizontal lines).

### Supplementary Methods

#### Isolation of high-molecular weight genomic DNA for PacBio sequencing

High-quality genomic DNA from argan (*Sideroxylon spinosum*) was isolated based on a modified CTAB protocol:

SOLUTIONS:

<u>Extraction buffer (EB):</u>
0.35 M sorbitol
100 mM Tris-HCl pH=7.5
10 mM EDTA

<u>Lysis Buffer (LB):</u>
200 mM Tris-HCl pH=7.5
50 mM EDTA
2M NaCl
2% w/v CTAB (Cetyltrimethylammonium bromide)

10% Sodium-N-lauroylsarcosinate

PROCEDURE:

- Grind ca 5g of frozen argan leaves to finest possible powder in mortar with N2liq.
- Resuspend in 30 ml EB in 50ml Falcon tube
- Spin at 10’000 rpm for 20 min in Falcon swingout rotor
- remove supernatant (SN)
- resuspend pellet in 5 ml lysis buffer; move VERY GENTLY with a sterile spatule
- Transfer to 15 ml Falcon tube and add 1% sodium-N-lauroylsarcosinate
- incubate at 65°C for 2h in water bath
- after cooling, add 2.5 ml chloroform:isoamylalcohol (=3-methyl-1-butanol) 15:1
- mix VERY GENTLY by mild rocking for several minutes
- Spin at 10’000 rpm for 30 min in Falcon swingout rotor
- Collect SN; avoid ANY traces of the interface (better to sacrifice part of the DNA!)
- add equal volume of isopropanol and mix VERY GENTLY. Whitish precipitate forms within minutes at room temperature. Move solution VERY GENTLY by slow rocking until precipitate forms a cloud
- transfer gDNA “cloud” with toothpick to a new 15 ml Falcon tube and dissolve in 3 ml of LB
- incubate at 65°C for 2h in water bath
- add 2 ml chloroform:isoamylalcohol 15:1
- mix VERY GENTLY by mild rocking for several minutes
- Spin at 10’000 rpm for 30 min in Falcon swingout rotor
- Collect SN in a new 15 ml Falcon tube; avoid ANY traces of the interface or lower phase
- add equal volume of isopropanol and mix VERY GENTLY by slow rocking
- after 30 min precipitation at RT collect gDNA (10’000 rpm for 20 min)
- wash pellet with 70% ethanol for 30 min; dry pellet for several hours
- Dissolve DNA in water / TE

#### Isolation of nuclei for HiC sequencing

Nuclei from argan (*Sideroxylon spinosum*) were isolated using the CelLyticTM Plant Nuclei Extraction Kit (Sigma™ Cat # CELLYTPN1) according to the protocol of the provider with minor modifications:

FINAL PROTOCOL:

Grind 4g leaves in N2liq.
Suspend in 20 ml 1x NIB (+1 mM DTT).
Filter through nylon mesh.
Spin down at 1260 g (3100 rpm) 10 min.
Resuspend pellet in 20 ml 1x NIB.
Add 1 % Triton-X-100, shake well.
Spin down at 12 g (300 rpm) 10 min.
Collect SN and shake; spin down at 50 g (600 rpm) 30 min.
Resuspend pellet in 7 ml NIB; add 1% Triton-X-100.
Load on gradient with 3 ml 2.3M sucrose / 3 ml 60% Percoll (**Figure S3**).
Spin at 3000 rpm (1270 g) for 30 min.
Collect Percoll phase (turbid; light green).
Dilute with 1x NIB and collect at 1500 rpm (320 g) for 30 min.
Wash with 1 ml NIB.
Spin at 14000 rpm for 10 min and suspend in 100 microL PURE nuclei buffer; snap freeze in N2

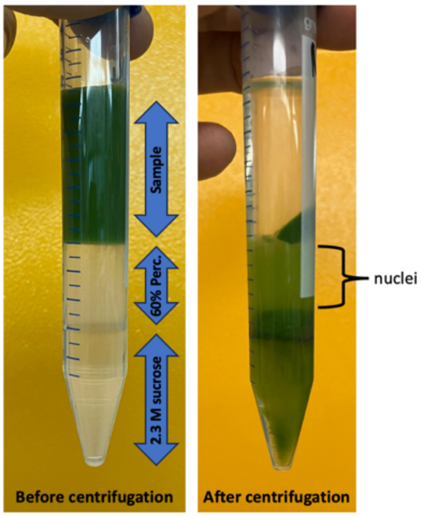


**Figure S3.** Isolation of argan nuclei by sucrose/Percoll gradient centrifugation

NIB: Nuclei Isolation Buffer

**Supplementary Table 1**

| Assembly | *S_spinosum*_hap1 | *S_spinosum*_hap2 | *S_spinosum*_QLOD |
|---|---|---|---|
| # contigs (>= 0 bp) | 14 | 20 | 186325 |
| # contigs (>= 1000 bp) | 14 | 20 | 79637 |
| # contigs (>= 5000 bp) | 14 | 20 | 8274 |
| # contigs (>= 10000 bp) | 14 | 20 | 2327 |
| # contigs (>= 25000 bp) | 14 | 20 | 268 |
| # contigs (>= 50000 bp) | 14 | 20 | 56 |
| Total length (>= 0 bp) | 636089469 | 654955621 | 690465237 |
| Total length (>= 1000 bp) | 636089469 | 654955621 | 617251183 |
| Total length (>= 5000 bp) | 636089469 | 654955621 | 478269350 |
| Total length (>= 10000 bp) | 636089469 | 654955621 | 438002149 |
| Total length (>= 25000 bp) | 636089469 | 654955621 | 408219428 |
| Total length (>= 50000 bp) | 636089469 | 654955621 | 401281704 |
| # contigs | 14 | 20 | 186325 |
| Largest contig | 116750276 | 121307699 | 49920449 |
| Total length | 636089469 | 654955621 | 690465237 |
| GC (%) | 34.09 | 34.18 | 32.57 |
| N50 | 50434390 | 50469191 | 24953461 |
| N75 | 43041221 | 42094784 | 3014 |
| L50 | 4 | 4 | 11 |
| L75 | 7 | 7 | 18670 |
| # N's per 100 kbp | 1.48 | 2.02 | 11831.95 |

**Supplementary Table 1**. Quast report of genome assemblies. The *S. spinosum* haplotig 1 and haplotig 2 assemblies were compared with a previous genome assembly of the species.

#### Supplementary Table 2

| | *S. spinosum* haplotig 1 | | | *S. spinosum* haplotig 2 | | |
|---|---|---|---|---|---|---|
| sequences | 14 | | | 20 | | |
| total length | 63608946 bp | | | 654955621 bp | | |
| GC level | 34.09% | | | 34.18% | | |
| bases masked | 398384631 bp ( 62.63 %) | | | 404898276 bp ( 61.82 %) | | |
| | number of elements* | length occupied | percentage of sequence | number of elements* | length occupied | percentage of sequence |
| Retroelements | 198079 | 154269473 bp | 24.25% | 196941 | 159838735 bp | 24.40% |
| SINEs: | 11075 | 1582809 bp | 0.25% | 6250 | 2922277 bp | 0.45% |
| Penelope: | 1620 | 354349 bp | 0.06% | 542 | 200823 bp | 0.03% |
| LINEs: | 47537 | 31611296 bp | 4.97% | 45128 | 25415231 bp | 3.88% |
| CRE/SLACS | 0 | 0 bp | 0.00% | 0 | 0 bp | 0.00% |
| L2/CR1/Rex | 0 | 0 bp | 0.00% | 0 | 0 bp | 0.00% |
| R1/LOA/Jockey | 74 | 84399 bp | 0.01% | 605 | 102268 bp | 0.02% |
| R2/R4/NeSL | 0 | 0 bp | 0.00% | 0 | 0 bp | 0.00% |
| RTE/Bov-B | 14374 | 8503015 bp | 1.34% | 13451 | 8435139 bp | 1.29% |
| L1/CIN4 | 25897 | 16294489 bp | 2.56% | 24235 | 14806276 bp | 2.26% |
| LTR elements: | 137847 | 120721019 bp | 18.98% | 145021 | 131300404 bp | 20.05% |
| BEL/Pao | 1952 | 344844 bp | 0.05% | 116 | 40209 bp | 0.01% |
| Ty1/Copia | 45741 | 47193634 bp | 7.42% | 46084 | 47782107 bp | 7.30% |
| Gypsy/DIRS1 | 52322 | 56896218 bp | 8.94% | 46424 | 53791999 bp | 8.21% |
| Retroviral | 2202 | 520755 bp | 0.08% | 2148 | 415671 bp | 0.06% |
| DNA transposons | 48486 | 18189792 bp | 2.86% | 37194 | 17378305 bp | 2.65% |
| hobo-Activator | 10701 | 5696725 bp | 0.90% | 10314 | 5358807 bp | 0.82% |
| Tc1-IS630-Pogo | 1106 | 546537 bp | 0.09% | 3028 | 1018803 bp | 0.16% |
| En-Spm | 0 | 0 bp | 0.00% | 0 | 0 bp | 0.00% |
| MULE-MuDR | 10146 | 5443525 bp | 0.86% | 4722 | 4043693 bp | 0.62% |
| PiggyBac | 0 | 0 bp | 0.00% | 0 | 0 bp | 0.00% |
| Tourist/Harbinger | 5671 | 1529072 bp | 0.24% | 2351 | 949529 bp | 0.14% |
| Other (Mirage, P-element, Transib) | 4688 | 701404 bp | 0.11% | 0 | 0 bp | 0.00% |
| Rolling-circles | 1848 | 1275553 bp | 0.20% | 3004 | 1396644 bp | 0.21% |
| Unclassified: | 615026 | 209454570 bp | 32.93% | 628986 | 210550985 bp | 32.15% |
| Total interspersed repeats | | 381913835 bp | 60.04% | | 387768025 bp | 59.21% |
| Small RNA: | 11988 | 6400808 bp | 1.01% | 7447 | 5968618 bp | 0.91% |
| Satellites: | 177 | 26404 bp | 0.00% | 2537 | 843040 bp | 0.13% |
| Simple repeats: | 238065 | 8394200 bp | 1.32% | 238722 | 8399224 bp | 1.28% |
| Low complexity: | 39289 | 1859318 bp | 0.29% | 39893 | 1897445 bp | 0.29% |

**Supplementary Table 2**. RepeatMasker report.

**Supplementary Table 3**

| *S. spinosum* haplotig 1. Results from dataset eudicots_odb10 | |
|---|---|
| C:97.8%[S:92.1%,D:5.7%],F:0.4%,M:1.8%,n:2326 | |
| Complete BUSCOs (C) | 2274 |
| Complete and single-copy BUSCOs (S) | 2142 |
| Complete and duplicated BUSCOs (D) | 132 |
| Fragmented BUSCOs (F) | 9 |
| Missing BUSCOs (M) | 43 |
| Total BUSCO groups searched | 2326 |

**Supplementary Table 3**. Busco output on *S. spinosum* haplotig 1 from dataset eudicots_odb10 containing 2326 BUSCOs.

**Supplementary Table 4**

| *S. spinosum* haplotig 2. Results from dataset eudicots_odb10 | |
|---|---|
| C:98.5%[S:92.6%,D:5.9%],F:0.3%,M:1.2%,n:2326 | |
| Complete BUSCOs (C) | 2291 |
| Complete and single-copy BUSCOs (S) | 2153 |
| Complete and duplicated BUSCOs (D) | 138 |
| Fragmented BUSCOs (F) | 7 |
| Missing BUSCOs (M) | 28 |
| Total BUSCO groups searched | 2326 |

**Supplementary Table 4**. Busco output on *S. spinosum* haplotig 2 from dataset eudicots_odb10 containing 2326 BUSCOs.
